# Spatio-temporal responses of butterflies to global warming on a Mediterranean island over two decades

**DOI:** 10.1101/2020.04.03.023689

**Authors:** Pau Colom, Anna Traveset, David Carreras, Constantí Stefanescu

## Abstract

In recent decades, efforts have been made to understand how global warming affects biodiversity and in this regard butterflies have emerged as a model group. The most conspicuous sign that warming is affecting the ecology of butterflies are the phenological advances occurring in many species. Moreover, rising temperatures are having a notable impact – both negative and positive – on population abundances. To date, patterns have generally been analysed at species level without taking into account possible differences between populations, which, when they are noted, are mostly attributed to large-scale climate differences across a latitudinal gradient. In this work, we use a long-term database of butterflies from the island of Menorca (Balearic Islands, Spain) to investigate how phenology and population dynamics have been affected by climate warming during the past two decades. In addition, we assess how responses are modulated by habitat characteristics and by species’ biological cycles. Our results show that species respond differently to warming at a local scale depending on season and habitat, and that coastal habitats in the Mediterranean region are particularly sensitive to the effects of climate change. Furthermore, our findings suggest that the effects of temperature could be partially offset in more inland habitats such as forests and deep ravines. The positive effect of temperature on ravine populations during the summer suggests that butterflies disperse across habitats as a response to rising temperatures during the season. This type of dispersal behaviour as a response to warming could be especially important in island ecosystems where the possibilities of modifying altitudinal or latitudinal distributions are often severely limited.

## Introduction

In recent decades, climate change has become a key factor in attempts to understand trends in biodiversity and in species’ distribution, phenology and population dynamics (Parmesan & Yohe 2003, Thomas *et al*., 2004, Araújo & Rahbek 2006, Bellard *et al*., 2012, Thackeray *et al*., 2016). A wealth of data has been assembled that shows how different organisms are responding to climate change, how populations are adapting to novel conditions and what the limits are to such adaptations (e.g. Devictor *et al*., 2012; Socolar *et al*., 2017; Radchuk *et al*., 2019). Of the model organisms on which much of this research has been focused, butterflies are ideal case studies for various reasons (Dennis, 1993; Parmesan, 2003). As ectothermic animals with short generation times, their development and activity are heavily constrained by environmental temperature; even so, rapid, noticeable responses at population level are commonly observed (e.g. Roy *et al*., 2001). Moreover, the existence of precise data on butterfly phenology (e.g. through standard monitoring methods; van Swaay *et al*., 1997; Schmucki *et al*., 2016) has afforded this insect group a key position in climate change research.

Advances in the flight periods of butterfly species related to increasing temperatures are well-established phenomena that have been reported from areas as diverse as the UK (Roy & Sparks, 2000), Central Europe (Altermatt, 2009) and the Mediterranean region (Stefanescu *et al*., 2003; Forister & Shapiro, 2003). Nevertheless, not all species respond to warming in the same way and different responses stemming from differences in species traits are known to occur. For instance, species with narrower dietary ranges in larval stages and more advanced overwintering stages have been found to exhibit greater advances in flight periods (Diamond *et al*., 2011). In addition, the relationship between butterfly phenology and temperature has been shown to be affected by habitat (Altermatt, 2012), elevation gradient (Gutiérrez-Illán *et al*., 2012), microclimate (WallisDevries & Van Swaay, 2006) and seasonality (Walther *et al*., 2002). Hence, phenological responses to warming will differ across populations of the same species.

The effects of warming on butterfly numbers are more difficult to predict as different factors may interact in complex ways in a context-dependent manner (e.g. Roy et al., 2001; WallisDeVries *et al*., 2011; Boggs and Inouye, 2012). Moreover, different climate conditions can have differing impacts on population dynamics depending on whether they act upon immature or adult stages (Radchuk *et al*., 2013).

In this work, we use a long-term database of butterflies from the island of Menorca (Balearic Islands, Spain) to investigate how the phenology and the population dynamics of this insect group have been affected by climate warming during the past two decades, and how their responses have been modulated by habitat characteristics and species’ biological cycles. As expected for a small island with no important mountain ranges (maximum elevation 358 m a.s.l.), the butterfly community is species poor and strongly dominated by common generalists. Nevertheless, most butterfly populations on the island have undergone similar declines to those reported from nearby continental areas where many more specialist species occur (Colom *et al.*, 2019). Climate warming has been suggested as a possible factor underlying these negative trends; recent work has revealed the major role of climate in the dynamics of Mediterranean butterflies and shown the negative impact of increasing temperatures and drought on certain species (e.g. Merrill *et al*., 2007; Zografou *et al*., 2014; Mills *et al*., 2017; Herrando *et al*., 2019; Carnicer *et al*., 2019). Moreover, given its small surface area, its physical limits as an island and the absence of high mountain ranges, species on Menorca are limited in their capacity to modify their distributions in response to climate change, as occurs in other areas (e.g. Parmesan *et al*., 1999; Wilson *et al*., 2005). Likewise, in contrast to mountainous areas (e.g. Gutiérrez-Illán *et al*., 2012), phenological variation between populations is expected to be weak or non-existent in Menorca due to its relatively flat landscape. In spite of this, systematic butterfly recording from Menorca for nearly two decades has revealed differences in phenology and population trends between monitored sites. Here we investigate in detail the nature of such variation and how it relates to topographical diversity on the island. The results are important for improving knowledge of how local conditions buffer the negative impacts of climate change in Mediterranean landscapes outside of mountain ranges, and provide guidelines as to which habitats are most deserving of concerted conservation efforts focused on mitigation and adjustment to climate change.

## Material and Methods

### Study area

Five sites on the island of Menorca were selected for study (Supplementary Figures S1 and S2), all close to the sea (1–10 km) and geographically separated by a minimum of 5 km from each other. They are at low altitude (0–50 m) and embrace in total 10 CORINE land-cover types (e.g. Mediterranean grassland, coastal marshland, coniferous Mediterranean forest and herbaceous crops). Although all five sites were used to define the overall phenology of the species on the island, we only used three with 15–18 years of data for comparisons: (1) Albufera des Grau, an open coastal area with dry meadows and Mediterranean scrub; (2) Santa Catalina, a forested area dominated by evergreen oak and pine forests; and (3) Barranc d’Algendar, a ravine in the south of the island with orchards at the beginning and thick undergrowth further inland. Although the climate is typically Mediterranean throughout the island, with hot-dry summers and mild-wet winters (Jansà et al., 2017), the geomorphology of its ravines generates moister microclimatic conditions that set Barranc d’Algendar apart from the other studied localities.

### Butterfly data

We used butterfly abundance data collected in 2001–2018 within the framework of the Catalan Butterfly Monitoring Scheme (CBMS) (www.catalanbms.org). Butterfly abundances were recorded weekly from March to September along fixed transects of 2–3 km in length using the standard BMS methodology (Pollard and Yates, 1993; Schmucki *et al*., 2016). Butterflies were counted and identified to species level in a 5×5 m area along transects (2.5 m to each side and 5 m in front of the recorder) whenever weather conditions met certain minimum standards (temperature over 15°C, preferably under a sunny sky).

Analyses focused on 16 of the 25 butterfly species recorded on Menorca that in all five localities have an annual frequency of occurrence of >0.5 and an annual mean abundance of over five individuals. After applying these criteria, we were left with data for 80 populations that had been monitored for 4–18 years. The 16 species selected are all habitat generalists (Colom *et al*., 2019) but still differed in certain ecological traits related to their voltinism (i.e. number of generations per year), migratory behaviour and hibernation phases, which were taken into account when analyzing phenological and population trends (Supplementary Table S1).

### Butterfly phenology

The flight periods of each species were identified and characterised after pooling abundance data for all five study sites for 2001–2018 (Supplementary Figure S3). This information, together with published data on the ecology of the species in Spain (García-Barros *et al.*, 2013; Vila *et al.*, 2018), allowed us to interpret the life cycle of these species on Menorca (Supplementary Table S1) and define periods on their flight curves that correspond to different generations (Table 1). Four species in our dataset are univoltine (*Callophrys rubi, Gonepteryx cleopatra, Maniola jurtina* and *Pyronia cecilia*), the remaining 12 being multivoltine. For each species, we established a *critical period* (CP, hereafter) into which the development of the immature stages – i.e. before adult emergence and after the diapause stage –was concentrated. We assumed that the temperature experienced during the CPs is crucial for accounting for annual variations in the flight period. According to knowledge of the biology of these species and a close examination of their flight curves on Menorca, we proposed a maximum of three possible variations in the CP for each species (i.e. a variable extended period before adult emergence) for which the mean temperature was calculated (Supplementary Table S2). For *G. cleopatra*, the CP in winter-spring does not coincide with the development of immature stages but with the time when hibernating adults are most sensitive to temperature changes as a prelude to breaking diapause.

**Table 1.** The Weighted Mean Flight Periods (WMFP) that were considered for each species and the critical periods that were selected in the best models. Significant relationships between WMFP and mean temperatures during the critical period (in the indicated region) are shown, with Beta negative values corresponding to advances of the WMFP with greater temperatures. Significant temporal trends in WMFP over the study period (2001–2018) are also shown (significant *P*-values are provided). WMFP correspond to spring (spr), early summer (esum), summer (sum), late summer (lsum), or to a succession of closely overlapping generations (over). *: univoltine species.

| Species | WMFP period<br>(weeks) | Climatic<br>region | Critical period | Temperature |  | Temporal |  |
| --- | --- | --- | --- | --- | --- | --- | --- |
|  |  |  |  | *P* | Beta | *P* | Beta |
| *Aricia cramera* | (11-26)sum | Menorca | May - July | 0.017 | -0.95 | - | - |
| *Callophrys rubi* * | (1-12)spr | Menorca | January - April | 0.023 | -0.71 | - | - |
| *Celastrina argiolus* | (1-14)spr | Menorca | March - April | 0.032 | -1.37 | 0.008 | 0.23 |
|  | (15-26)sum | - | - | - | - | - | - |
| *Coenonympha pamphilus* | (1-13)spr | Menorca | February - April | 0.026 | -0.59 | 0.04 | -0.08 |
|  | (14-26)sum | Menorca |  | - | - | - | - |
| *Colias crocea* | (1-10)spr | - | - | - | - | - | - |
|  | (11-20)sum | Menorca | May - July | <0.0001 | -0.78 | - | - |
| *Gonepteryx cleopatra* * | (1 - 9)spr | - | - | - | - | - | - |
|  | (11 - 20)sum | Menorca | April - June | 0.0004 | -3.74 | - | - |
| *Lasiommata megera* | (1 - 9)spr | - | - | - | - | - | - |
|  | (10 - 22)sum | - | - | - | - | - | - |
| *Lycaena phlaeas* | (1 - 15)spr | - | - | - | - | - | - |
|  | (16 - 30)sum | - | - | - | - | - | - |
| *Maniola jurtina* * | (1-17)spr | Menorca | March - June | 0.011 | -0.31 | - | - |
|  | (18 - 30)sum | - | - | - | - | - | - |
| *Pararge aegeria* | (1 - 30)over | Menorca | May - July | 0.008 | -1.02 | - | - |
| *Pieris brassicae* | (1 - 22)spr-<br>esum | North Africa | March - April | 0.016 | -2.49 | - | - |
|  | (23 - 30)lsum | Menorca | July - September | 0.041 | -0.57 | 0.003 | 0.13 |
| *Pieris rapae* | (1 - 30)over | North Africa | April - July | 0.042 | -0.89 |  |  |
| *Polyommatus celina* | (1 - 9)spr | - | - | - | - | - | - |
|  | (11 - 24)sum | Menorca | May - August | <0.0001 | -1.03 | - | - |
| *Pyronia cecilia* * | (10 - 30)sum | Menorca | May -June | 0.001 | -0.68 | - | - |
| *Vanessa atalanta* | (1 - 17)spr | - | - | - | - | - | - |
|  | (18 - 24)sum | - | - | - | - | - | - |
| *Vanessa cardui* | (1 - 10)spr | North Africa | January - April | 0.044 | -0.96 |  |  |
|  | (11 - 20)esum | - | - | - | - | - | - |
|  | (21 - 30)lsum | Med. Europe | July - September | 0.007 | -1.17 | - | - |

To estimate the annual flight period of a butterfly population, we used the Weighted Mean Flight Period (hereafter WMFP), which is a statistic widely used in butterfly phenological studies (e.g. Roy & Sparks, 2000; Stefanescu *et al*., 2003: Gutierrez-Illán *et al*., 2012). The WMFP represents the date (i.e. the week in a range of 1–30) in which the mean of adult counts of a given species at a given locality occurs:

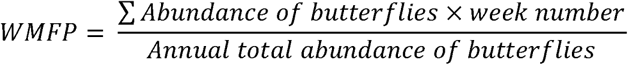

As in other studies (Pollard *et al*., 1991, Stefanescu *et al*., 2003), the recording weeks rather than the day of the counts were used as the time unit. For multivoltine species, we calculated two or three WMFPs corresponding to spring and summer (in some cases early-summer or late-summer) generations (see Table 1 for the WMFPs calculated for each species). As an exception, we calculated a single WMFP for two multivoltine species with overlapping generations (*Pararge aegeria* and *Pieris rapae*) given that their overall flight curves were better rendered by a unimodal pattern.

### Temperature data

Detailed climatic records for Menorca are available for the last 48 years from various meteorological stations run by the Spanish Meteorological Agency (AEMET). During this period, the mean temperature on Menorca has increased 0.34°C/decade (0.41°C/decade for maximum temperatures). This increase, however, was not uniform over the seasons as the late-spring months (April, May and June) were the greatest contributors to the overall annual warming (Jansà & Gomis, 2018).

To investigate the relationship between temperature and phenology, we associated butterfly recording sites with the closest meteorological station with data available for 2001–2018. At each site, the WMFP of a given species was then related to the mean temperatures of the months covering its critical period of development (Supplementary Table S2).

Given that some of the butterflies on Menorca are migratory species (e.g. *Colias crocea, Pieris rapae, Pieris brassicae, Vanessa atalanta* and *Vanessa cardui*) whose immature stages do not develop on the island, it was also necessary to obtain the mean temperatures in the regions where these species originate. Specifically, through ERA-interim, a global atmospheric reanalysis updated in real time by the European Centre for Medium-Range Weather Forecasts (www.ecmwf.int), the monthly mean temperatures of four domains were obtained: North Africa (32.25°N < latitude < 36.75N°; 10.50°W < longitude < 10.50°E); Europe (32.25°N < latitude < 45.75°N; 10.50°W < longitude < 10.50°E); Mediterranean Europe (37.50°N < latitude < 45.75°N; 0.00° < longitude < 10.50°E); and Atlantic Europe (37.50°N < latitude < 45.75°N; 10.50°W < longitude < 0.75°W).

### Statistical analyses

#### Phenological analysis

We used Generalised Linear Mixed Models (GLMM) to test whether or not species advanced their phenologies in warmer years (i.e. warmer critical periods) and whether or not there were any related trends during the study period (2001–2018). Tests were conducted for the different flight periods of each species (except for *Pararge aegeria* and *Pieris rapae*, for which it was impossible to distinguish between individual generations due to overlaps; see above). The mean temperatures in the various combinations of the critical months (as defined previously) were used as the explanatory variable in the different models, with ‘site’ used as a random factor (Supplementary Table S2). The best models were selected based on the lowest AIC value (Akaike Information Criterion).

We used the results of these first analyses to examine with linear regressions the relationship between the WMFPs and the mean temperature of the relevant CPs. The strength of the relationship was quantified through the beta coefficient of the regression and was compared between localities (the three localities with >15 years of data) and seasonality (spring vs. summer generations) using a GLMM in which ‘site’ and ‘season’ were set as fixed factors and ‘species’ as a random factor.

#### Population dynamics analysis

We followed a similar approach to study the effects of temperature on population abundances in different generations. We examined separately the effect of temperature on immature stages and on adults by (1) using the mean temperatures of the CPs (for immatures) that were selected in the best models in the first analysis, and (2) using the mean temperatures of the flight time period in which butterflies were recorded in each population. In both cases, we analysed the differences between localities and seasonality (i.e. spring vs. summer generations). To compare temperature effects between localities and between seasons, we excluded the long-distance migrants *V. cardui* and *P. brassicae* given that the phenology and abundance of their populations are not related to local temperatures (see Table 1).

## Results

### Temporal trends in temperature

In spite of the general warming trend in Menorca since the 1970s, we did not detect any overall significant trend in mean annual temperatures in the study period (2001–2018). The only month with a marginally significant increase in mean temperature was April (*P* = 0.084, *R*^*2*^ = 0.18). Nevertheless, we did find a significant annual increase of 0.78 °C/decade in maximum temperatures (*P* = 0.0009, *R*^*2*^ = 0.51). Moreover, when we restricted the analysis to individual months, there were significant warming trends in April (*P* = 0.003, *R*^*2*^ = 0.444), May (*P* = 0.066, *R*^*2*^ = 0.15), September (*P* = 0.009, *R*^*2*^ = 0.35) and December (*P* = 0.007, *R*^*2*^ = 0.38) (Fig. 1).

**Figure 1.**
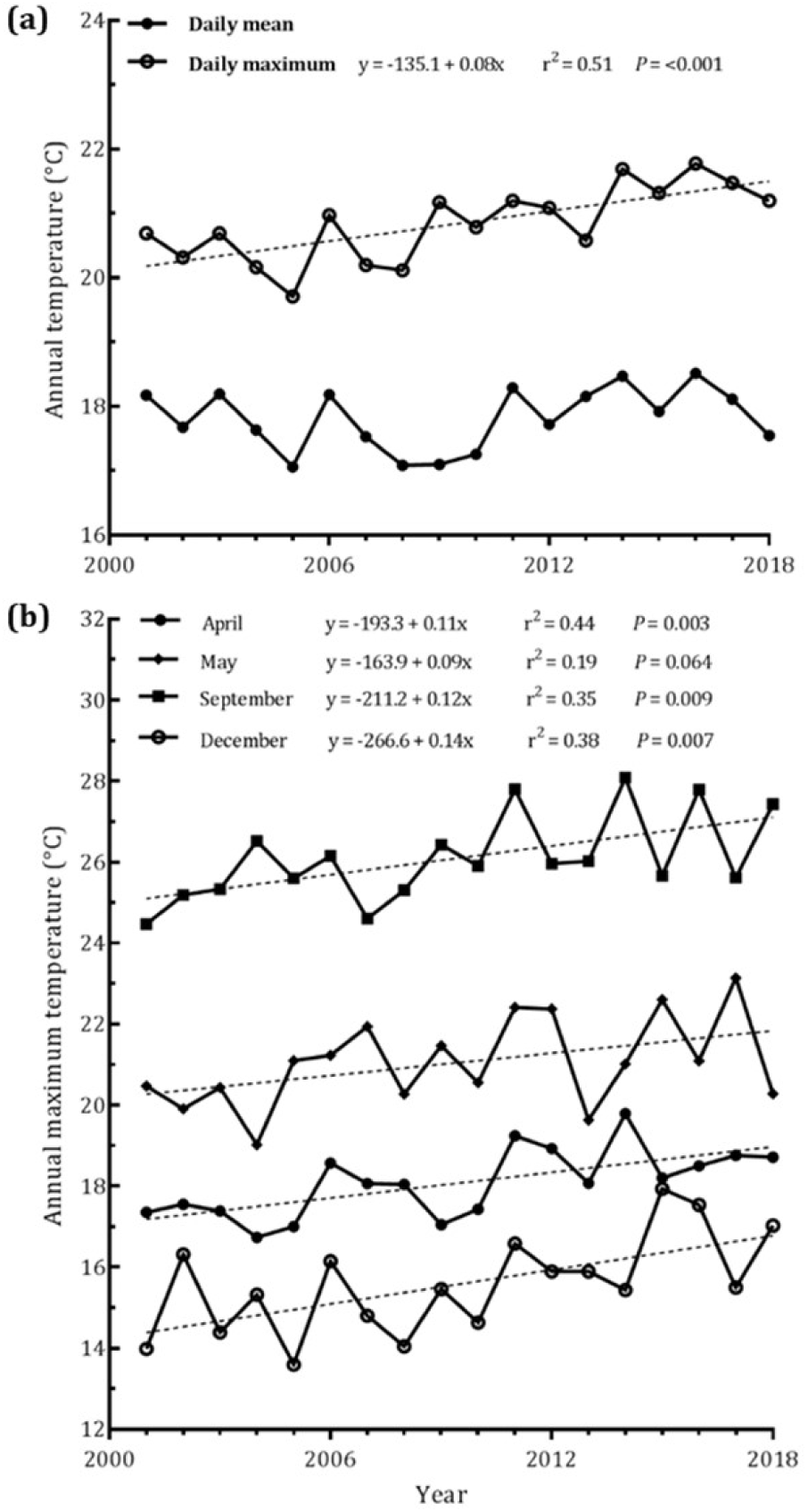
Climate trends in Menorca in 2001–2018. (a) Mean and maximum annual temperatures. (b) Mean monthly maximum temperatures from April, May, September and December. Dotted lines represent significant or marginally significant linear regressions (*P* < 0.1).

### Phenological response to increasing temperatures

Three species changed the timing of their WMFPs during the study period (Table 1): *Coenonympha pamphilus* advanced the WMFP of its spring generation (*P* = 0.04), while there were significant delays in the spring WMFP of *Celastrina argiolus* (*P* = 0.008) and in the late summer WMFP of *Pieris brassicae* (*P* = 0.03). Despite these temporal delays, in both *C. argiolus* and *P. brassicae* there was a significant negative relationship between their WMFPs with temperature (Table 1).

In all, 80% of the analysed species showed the same negative WMFP-temperature relationship in some of their generations (Table 1). In fact, only three species (*Lasiommata megera, Vanessa atalanta* and *Lycaena phlaeas*) did not advance their phenology with increasing temperatures.

In multivoltine species with clearly distinct flight periods (*A. cramera, C. argiolus, C. pamphilus, C. crocea, P. brassicae, P. celina* and *V. cardui*), phenological advances were recorded in both spring and summer (four and five cases, respectively; Table 1). In the two long-distance migrants *V. cardui* and *P. brassicae*, WMFPs did not have significant relationships with local temperatures but with mainly temperatures in the presumed regions of origin of the recorded adults (i.e. North Africa for the spring generations of both species, and Mediterranean Europe for the late summer generation of *V. cardui*).

For two species, *Pararge aegeria* and *Pieris rapae*, the great overlap in their generations did not allow us to analyse their spring and summer phenologies separately. Nevertheless, the global unimodal flight curves of both species (weeks 1–30) showed a significant negative relationship with late-spring–early-summer temperatures (April–July; Table 1).

The four univoltine species in our dataset also showed phenological advances with increasing temperatures. As before, in the two species with unimodal flight curves, advances were recorded in spring (*Callophrys rubi*) and in summer (*Pyronia cecilia*) (Table 1). Despite having a single annual generation, the flight curves of the other two univoltine species were approximately bimodal. In the case of *G. cleopatra*, a first small peak corresponds to adults from the previous season that come out of hibernation in early spring, while a second and stronger peak corresponds to the new annual generation. We found a significant negative WMFP-temperature relationship in this second flight period. In *M. jurtina*, there was a strong first peak in late spring that corresponds to the emergence of the annual generation, followed by a much more prolonged but blurred second peak that mainly consists of the females that survive through the summer but do not begin oviposition until September. Only the first flight period of this species showed a significant relationship with temperature (Table 1).

### Spatial and seasonal variation of the WMFP-temperature relationship

The WMFP-temperature relationship was explained both by the seasonal (i.e. spring and summer flying periods) and spatial (i.e. the three individual sites) variables. For the 14 species analysed, temperature had a greater negative effect in spring than in summer (−0.65 ± 0.86 vs. - 0.34 ± 0.84; *P* = 0.0478) (Fig. 2). The analysis also showed marginally significant differences between localities (*P* = 0.081). Specifically, there was a weaker WMFP-temperature relationship in Barranc d’Algendar (i.e. the ravine) than in the other two areas, Albufera des Grau (coastal area, *P* = 0.0471) and Santa Catalina (Mediterranean forest, P = 0.0651) (Fig. 2). The latter two populations did not differ significantly.

**Figure 2.**
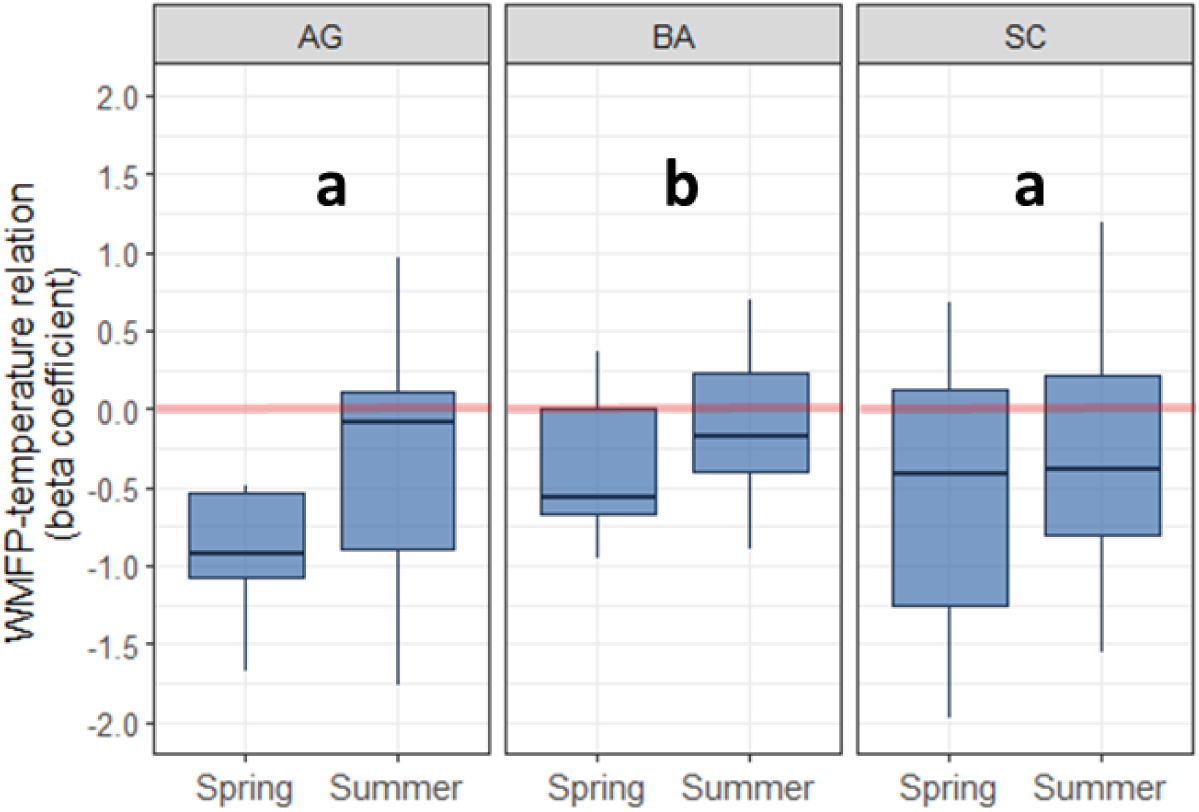
Beta coefficient of the regression of the Weighted Mean Flight Period (WMFP) and temperature at different localities (*P* = 0.081) (AG: Albufera des Grau: coastal area; BA: Barranc d’Algendar: ravine area; SC: Santa Catalina: Mediterranean forest) and seasons *(P* = 0.048) (spring and summer). Letters show significant differences between localities.

### Spatial and seasonal variation in the abundance-temperature relationship

The analysis of population abundance showed similar results for the two types of models, i.e. considering the temperatures during the CP before adult emergence and during the butterfly’s flight period. Both models showed significant differences between localities (model 1: *P* = 0.001; model 2: *P* = 0.058) and between seasons (model 1: *P* = 0.09; model 2: *P* = 0.005). However, in the first model (i.e. for developmental periods) the spatial variable was more important than the seasonal variable, whereas in the second model (i.e. for adult flight periods) the reverse was true.

As for the spatial scale, it was clear from both models that butterfly populations in the ravine (Barranc d’Algendar) benefitted more from higher temperatures than in the two other sites (Fig. 3). This was particularly the case in summer, when higher temperatures led to more abundant butterfly populations in the ravine (beta coefficients were positive in both models) but gave rise to less abundant populations in the coastal and forest sites (beta coefficients were negative at both sites and in both models).

**Figure 3.**
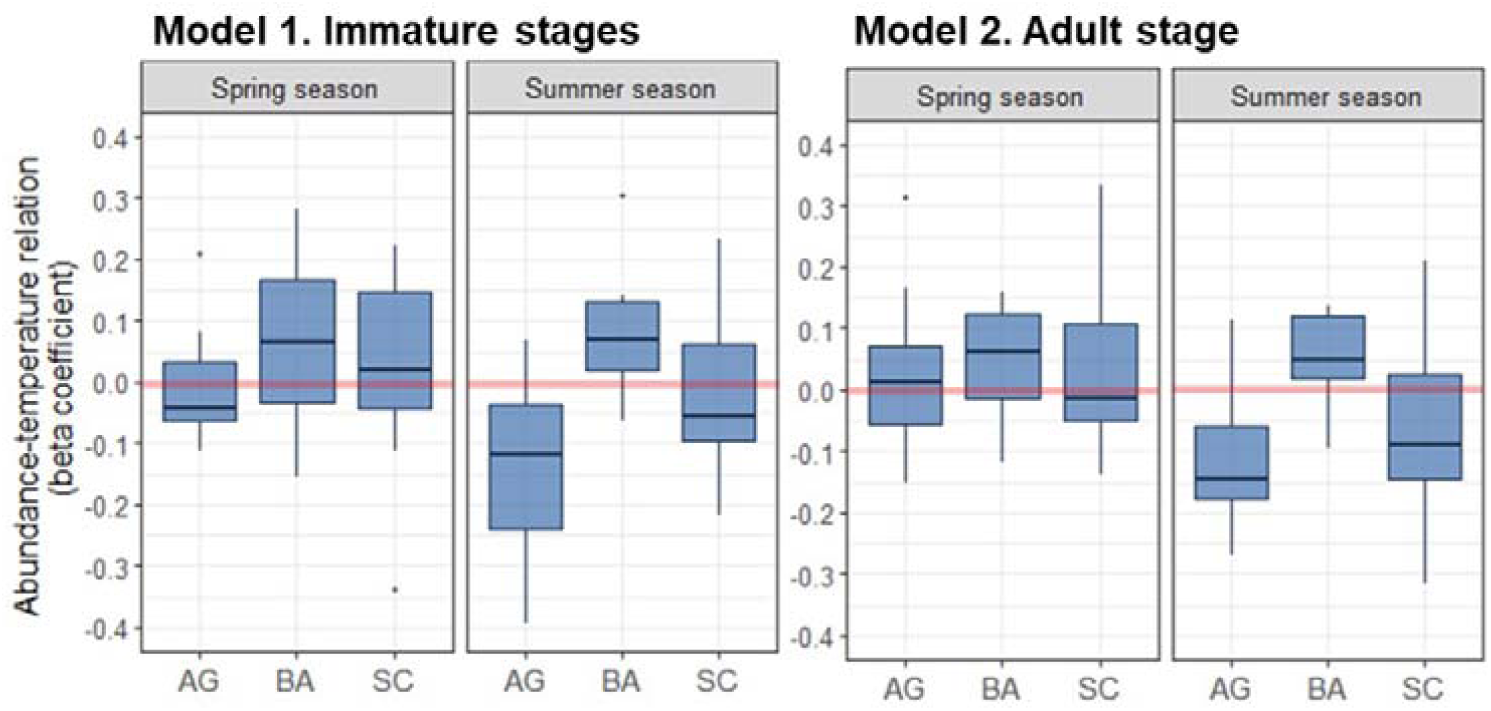
Beta coefficient of the regression of butterfly abundance and temperature at different localities (AG: Albufera des Grau: coastal area; BA: Barranc d’Algendar: ravine area; SC: Santa Catalina: Mediterranean forest) and seasons (spring and summer). Model 1 represents the effect of temperature on the critical periods of the species (i.e. the immature stages). Model 2 represents the effect of temperature on the adult stages. Both models show significant differences between seasons ((1) *P* = 0.09; (2) *P* = 0.005) and localities ((1) *P* = 0.001; (2) *P* = 0.058).

We found a consistent stronger temperature effect on population abundance in summer than in spring (Fig. 3). This was especially so at the coastal and forest sites (differences in beta coefficients between spring vs. summer: model 1 = 0.04; model 2 = 0.03) compared to the ravine site (differences in beta coefficients between spring vs. summer = model 1 = 0.02; model 2 = 0.009). Interestingly enough, in both the coastal and forest areas there was a reverse effect of the temperature on population abundance, with an almost neutral effect in spring but a very strong negative effect in summer. In the ravine area, by contrast, the effect was always positive and very similar in spring and summer.

## Discussion

Global warming is not spatially homogeneous and its effects vary widely across biogeographical areas (Post *et al*., 2018), the Mediterranean region, in particular, being one of the most sensitive areas to this phenomenon (Giorgi, 2006). Likewise, warming is not homogeneous throughout the year and in the Mediterranean region the greatest contribution to annual warming occurs in the transition from spring to summer (Garcia, 2015). There is also strong evidence to suggest that summer seasons in the Mediterranean are becoming longer (Bartolini *et al*., 2012; Jansà *et al*., 2017). In this sense, despite the short climatic period analysed in our study, warming trends (with respect to maximum temperatures) were observed in April, September and December, and also marginally in May. Moreover, the annual increase in maximum temperature in the study period (0.78°C/decade) is higher than the annual increase reported recently for Menorca in the period 1971–2016 (0.41°C / decade) (Jansà & Gomis, 2018). This trend may be related to the increase in the frequency of extreme climatic events (i.e. unusually high temperatures) observed in other parts of the Mediterranean, especially in the warm season (Caloiero *et al*., 2017; Scorzini *et al*., 2018).

Despite the observed warming trends during the study period, however, we only detected phenological trends in three species. On the one hand, *Coenonympha pamphilus* advanced its phenology in spring, a finding consistent with other studies of this species in the UK (Roy & Sparks, 2000) and the continental Mediterranean region (Stefanescu *et al*., 2003), whilst, on the other, *Celastrina argiolus* and *Pieris brassicae* showed phenological changes in the opposite direction, i.e. delayed phenologies. In *C. argiolus*, the apparent delay in their spring WMFP is probably an artefact related to an advance in the summer generation, which has many more individuals and may overlap with the spring generation in warmer years. For *P. brassicae*, the delay in the summer WMFP may have different causes. Firstly, this species has a complex life cycle in the Mediterranean region where it enters a summer diapause or aestivation phase that is not observed in other regions, as noted by Held & Spieth (1999) and Spieth *et al*. (2011); according to these authors, more severe drought conditions could lengthen the summer diapause period, resulting in a later emergence of the summer generation. Moreover, *P. brassicae* is migratory and many specimens move to higher latitudes to avoid the dry summer season (CS, pers. obs.). Conversely, in late summer, large numbers of *P. brassicae* from Central Europe migrate southward to breed in the Mediterranean region. The arrival of these specimens on Menorca, which in some cases is massive, could be delayed due to an extension of the summer season given that higher temperatures allow for an increase in the number of generations developed in Central Europe (Altermatt, 2009).

Overall, our findings are consistent with the idea that warming causes significant advances in the flight period of most butterflies. The only three species for which this pattern was not recorded are rare at some of the monitored localities and, therefore, their very low abundances probably precluded the detection of the expected advancement.

As hypothesized, we found evidence that the phenology of the long-distance migrants *Pieris brassicae* and *Vanessa cardui* is not influenced by local temperatures but, rather, by temperatures experienced in distant source areas. Interestingly, for the rest of the species affected by local temperatures we found that phenological patterns were also strongly dependent on both habitat and season, which implies that variation occurs between populations of the same species (Figs. 2, 3). On a temporal scale, the effects of temperature were not seasonally homogenous and, although advances did occur in both spring and summer, warming caused the strongest effects in the spring generations. This pattern, observed in numerous other studies of different groups of organisms (e.g. Walther *et al.*, 2002), is a consequence in the case of butterflies of how the metabolism of immature stages relates to environmental temperatures. It is well known that in insects the rate of development depends on body temperature, which in turn, as ectothermic organisms, is closely contingent on external temperature. Once a threshold of minimum temperature is exceeded, developmental rates increase linearly for a range of temperature until a decline occurs as an upper threshold is approached, beyond which no further development is possible (cf. Kingsolver, 1989). In the Mediterranean region, this means that temperature increases in the winter-spring transition, when average temperatures are commonly around minimum thresholds, provoke sharper developmental increases than when they occur in summer, when temperatures may even reach the upper developmental thresholds. Moreover, in the current context of an increase in the frequency of extreme climatic events (Easterling *et al*., 2000; Coumou & Rahmstorf, 2012), greater fluctuations in temperature (even if the average is constant) may mean a decrease in the developmental rate in warm conditions but an opposite effect at low temperatures (Paaijans *et al*., 2013).

Our results also suggest a seasonal pattern for the effects of temperature on abundance (Fig. 3). While in the spring the effects of higher temperatures were very variable during both the development of immature stages and the adult flight period, in summer they clearly had a negative effect, except in the ravine. During summer, the increase in temperature generally implies an increase in aridity and a reduction in water availability, which has been identified as a key limiting factor for butterfly species in the Mediterranean (Stefanescu *et al*., 2011; Herrando *et al*., 2019) and for plant-butterfly interactions (Donoso *et al*., 2016). In fact, the difference between this negative effect in summer and the neutral effect in spring was more noticeable when we considered the adult flight period (Fig. 3, model 2). During the summer season, a decrease in nectar supply under drought conditions could explain the negative effect of temperature on butterfly abundances (WallisDeVries *et al*., 2012). On the other hand, in spring, which is the rainy season in the study region, temperature increases are not as strongly associated with drought. In spring, however, multiple causes could explain a negative or positive effect of temperature on the abundance of the first butterfly generations. On the one hand, faster development due to higher environmental temperatures could lessen mortality during larval and pupal stages as the exposure time to their predators and parasitoids is reduced (e.g. Pollard, 1979). But, on the other hand, an earlier emergence of butterflies is not necessarily accompanied by an advance in plant flowering. Consequently, asynchrony could occur between butterfly and plant phenologies, especially in dry springs (Donoso *et al*., 2016), with a negative effect on butterfly abundances.

On a spatial scale, differences in phenology between populations of the same species have usually been attributed to adaptive or plastic responses to local day-length and temperatures (Hodgson *et al*., 2011). However, for nearby populations with no marked altitudinal gradient, these factors are minimized. In spite of this, our results showed significant differences in the effect of temperature on both the flight period and the abundance of different populations. Our results agree with those reported by Altermatt (2012), who showed that habitat (in his study due to anthropogenic changes in microclimate) can affect butterfly phenology. In our case, the strongest effects on phenology and abundance were observed at the coastal site. This suggests that in warmer years the effect of temperature could be more notable on coastal than in forested areas, where more stable climatic conditions are expected (cf. Suggitt *et al*., 2011).

The case of Barranc d’Algendar area deserves special mention. The butterfly response to warming at this site is evidently different from that at the other two localities. The less marked effect of temperature on the phenology of ravine populations and the increase in their abundance in warmer summers could be explained by the particular environmental conditions of these habitats. The milder temperatures in summer in this and other ravines could favour the arrival of individuals of highly mobile species from other more arid environments on the island since in the ravines there is a greater abundance of both adult (i.e. nectar supply) and larval resources (i.e. host plants) due to greater humidity. On an island like Menorca, where species have a very narrow latitudinal and altitudinal range in which to respond to warming, dispersal into such environments may be a common response. The geomorphology of the ravines in Menorca could lead to a beneficial microclimate for butterfly species during certain critical periods, especially under extreme weather conditions. Consequently, the ravines constitute biodiversity reservoirs in Menorca due to their capacity to act as a buffer for extreme weather conditions.

## Acknowledgements

We would like to thank the Observatori Socioambiental de Menorca, S’Albufera des Grau Natural Park and the Agencia Reserva de la Biosfera de Menorca for their support for the recording stations. PC is funded by a PhD fellowship financed by the Govern de les Illes Balears (FPI-CAIB-2018), part of project CGL2017-88122-P and financed by the Spanish Government. The Catalan Butterfly Monitoring Scheme is funded by the Departament de Territori i Sostenibilitat de la Generalitat de Catalunya. Michael Lockwood revised the Engish version.

## Supporting Information

### Figures

**Figure S1.**
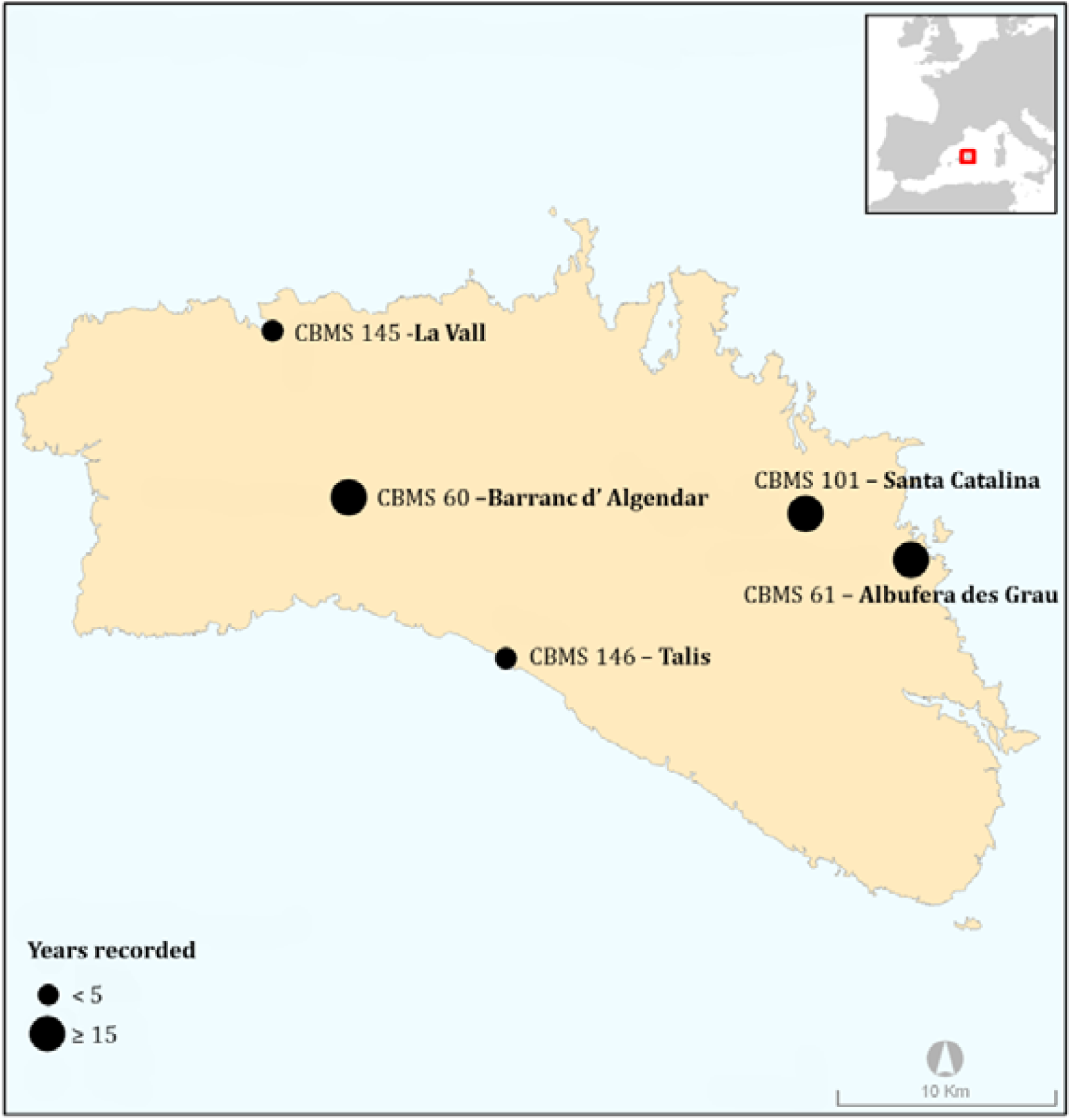
Map of the study area. The five sites form part of the CBMS network (www.catalanbms.org) and were used to characterize the phenology of the species. However, only the three sites with over 15 years of data were used in the analyses.

**Figure S2.**
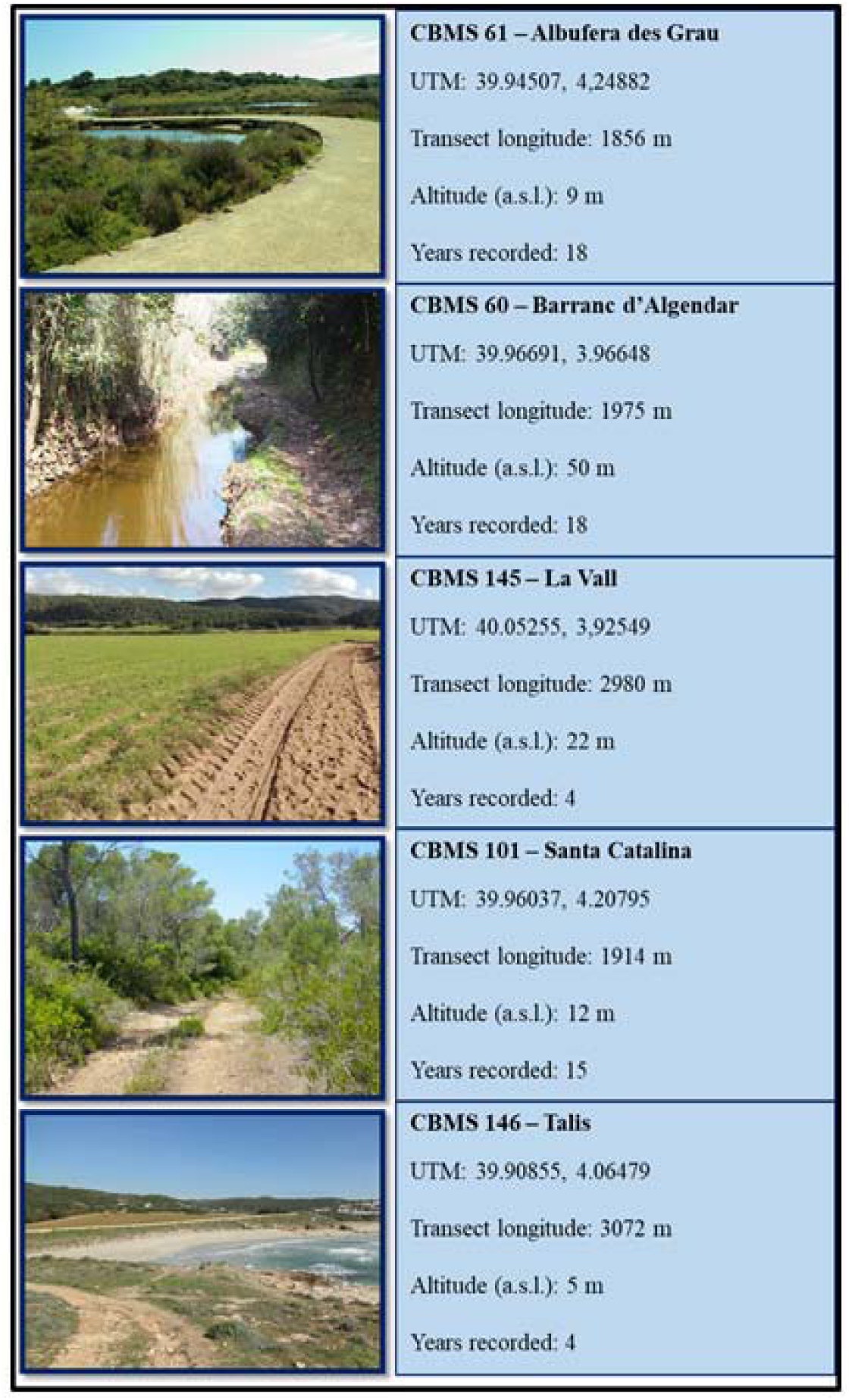
Details of the five CBMS transects used in the study.

**Figure S3.**
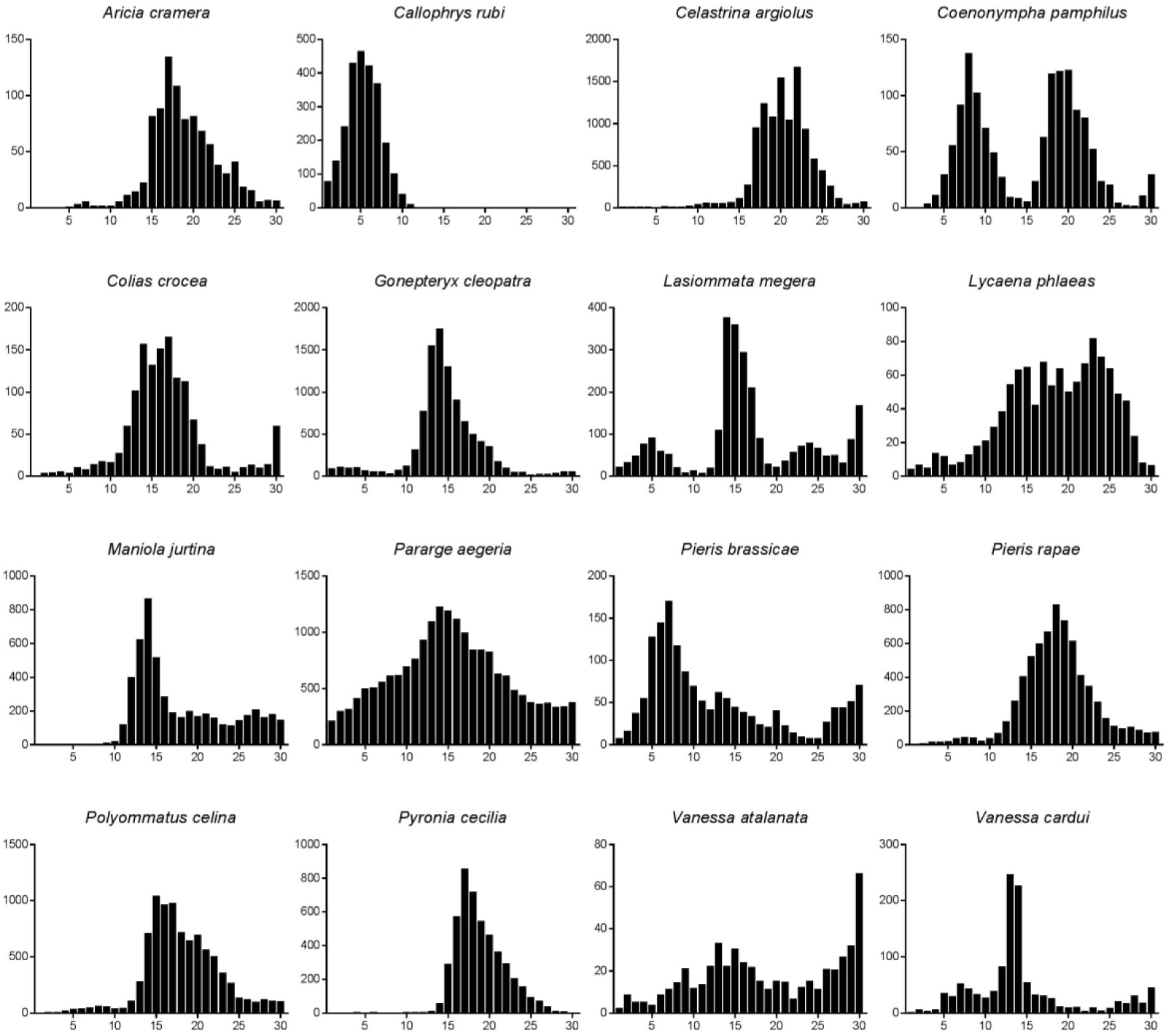
Butterfly species flight curves. The x-axis represents the 30 weeks of monitoring running from the first week of March to the fourth week of September. The y-axis represents the mean abundance standardized to 100 m of transect length.

### Tables

**Table S1.**
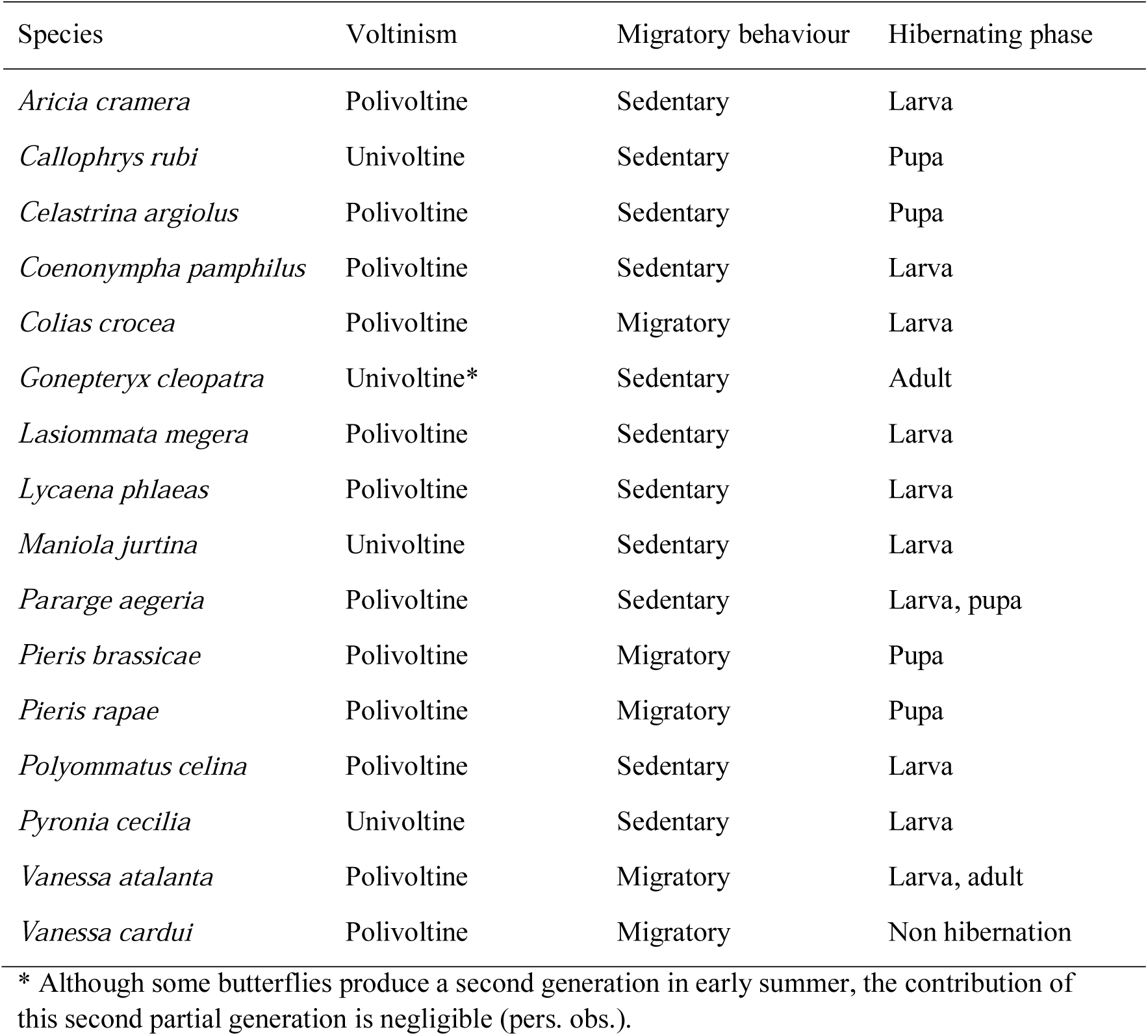
Ecological traits of the analysed butterfly species.

**Table S2.**
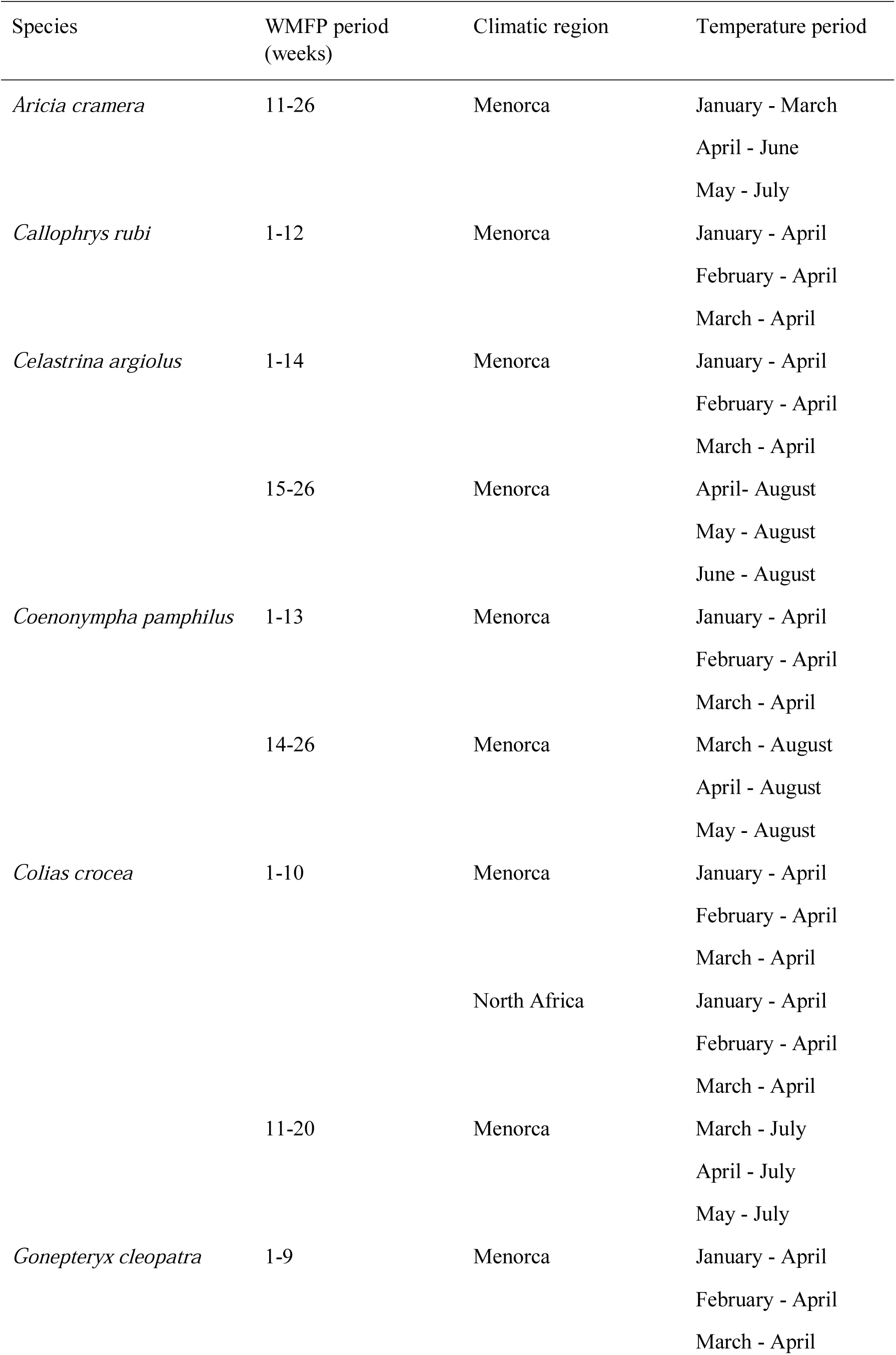
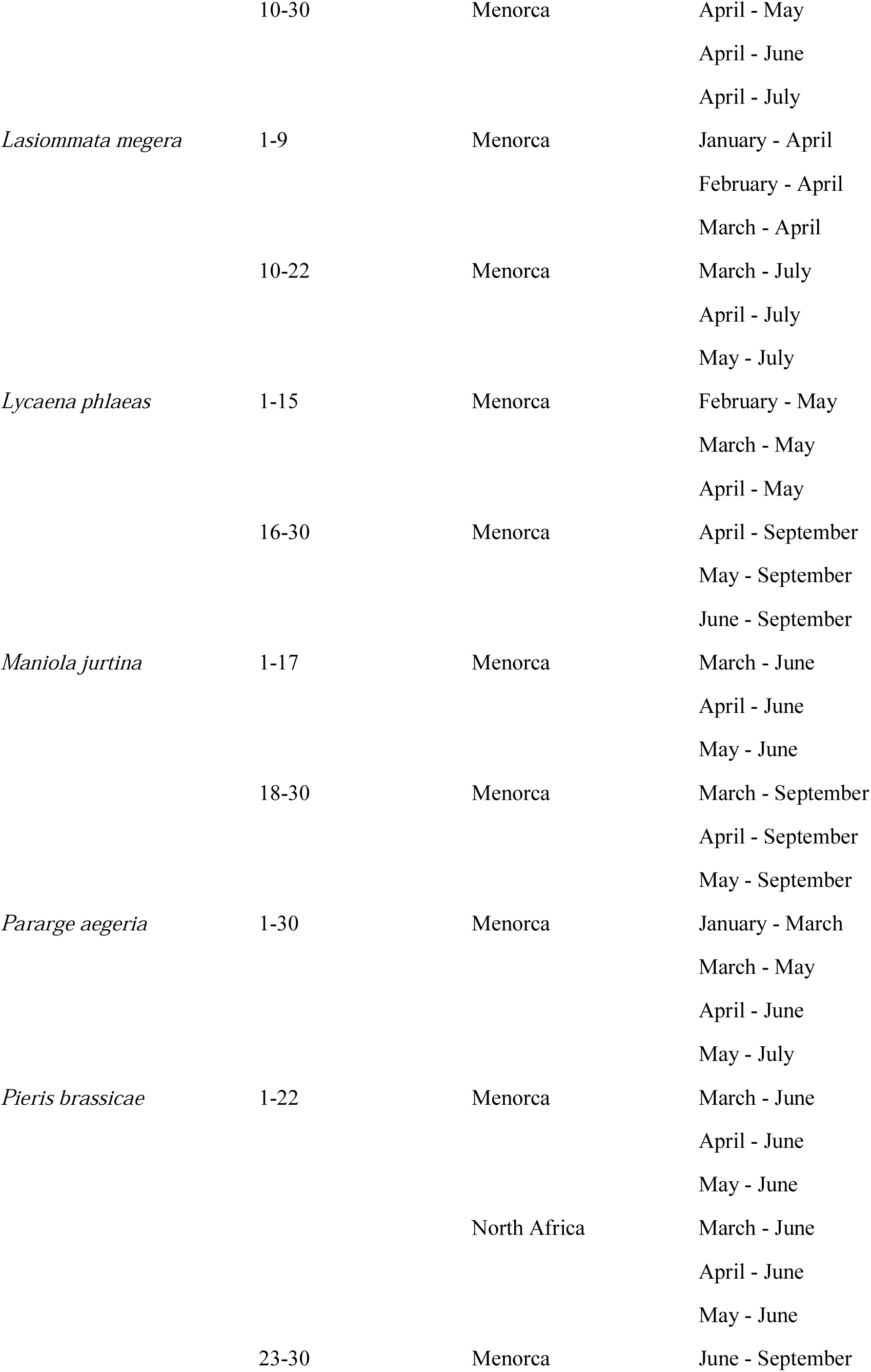
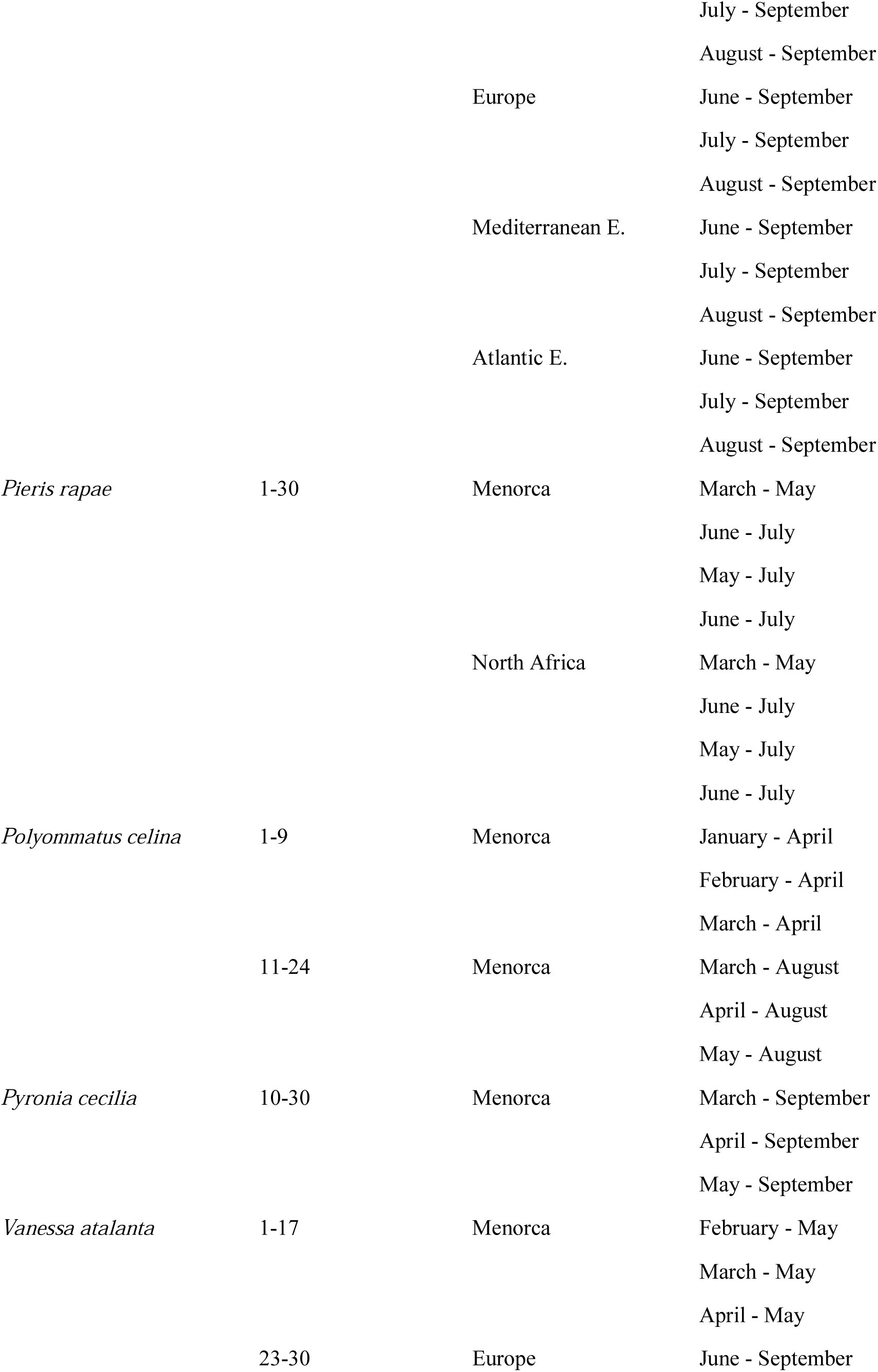
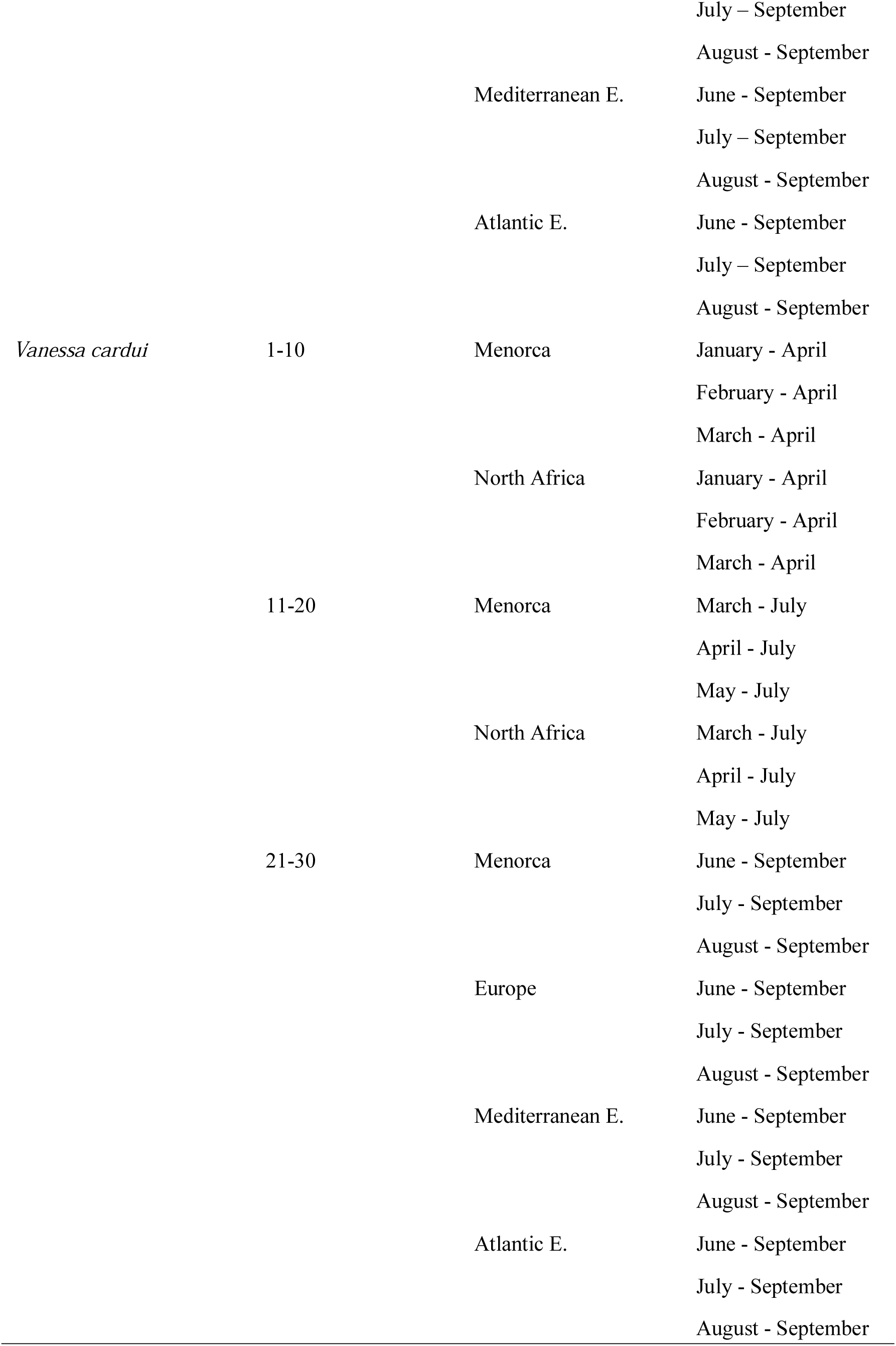
Different combinations of months and climatic regions used to analyse the critical periods of butterfly species in both spring and summer.

